# Identification and classification of innexin gene transcripts in the central nervous system of the terrestrial slug *Limax valentianus*

**DOI:** 10.1101/2020.12.21.423878

**Authors:** Hisayo Sadamoto, Hironobu Takahashi, Suguru Kobayashi, Hirooki Kondoh, Hiroshi Tokumaru

## Abstract

In invertebrates, innexin is involved in the formation of single-cell membrane channels and intercellular gap junction channels. Generally, there are multiple isoforms of innexin family proteins in various animal species, which enable the precise regulation of channel function. In molluscan species, the sequence information of innexins is still limited and the sequences have not been classified.

This study examined the innexin transcripts expressed in the central nervous system of the terrestrial slug *Limax valentianus* and identified 16 transcripts of 12 innexin isoforms, including the splicing variants. To examine the function of molluscan innexin isoforms, phylogenetic analysis was performed using the innexin sequences of molluscan species. Next, the phosphorylation, N-glycosylation, and S-nitrosylation sites in the isoforms were predicted to characterize the innexin isoforms. Further, 16 circular RNA sequences of nine innexin isoforms were identified in the central nervous system of *Limax*. The identification and classification of the gene transcripts of molluscan innexins provided novel insights for understanding the regulatory mechanism of innexins in the central nervous system.

## Introduction

Gap junction channels are formed by docking of single-cell membrane channels between adjacent cells, and that allow intercellular communication via the exchange of small molecules such as ions, nucleotides, small peptides and micro RNA (miRNA). Recent studies have indicated that the gap junction-related proteins are involved in the regulation of brain function. Neuronal gap junctions, which are known as electrical synapses, exhibit plasticity (1)(2). The single-cell membrane channels are involved in neuroplasticity through interaction with the chemical synapses (3)(2). The channel properties of gap junction-related proteins are regulated through protein modifications, such as N-glycosylation, S-nitrosylation, and phosphorylation (4)(5)(6)(7).

Three family proteins, innexin in invertebrate, connexin and pannexin in vertebrate, are known as the gap junction-related proteins. These three family proteins share structural features that are characterized by four transmembrane domains, two extracellular loops, and intracellular N- and C-terminal domains (8). The hexamers, heptamers, and octamers of these proteins form a single-cell membrane channel (9)(10). Additionally, the multiple isoforms of these proteins contribute to the specialized regulation of the channel property and the intercellular docking of the single-cell membrane channels. Previous studies have also reported the differential characteristics of the three family proteins (11)(12). In vertebrates, connexin proteins form both single-cell membrane channels and intercellular gap junctions. However, pannexin proteins form only single-cell membrane channels *in vivo* as glycosylation at their extracellular domain inhibits the formation of intercellular gap junction channels. In invertebrates, innexin exhibits the functions of both connexin and pannexin. The classification and function of invertebrate innexin isoforms have not been completely understood.

In molluscan species, molecular biological analyses of innexins remain even untouched, even though various studies have examined the electrical synapses in the central nervous system (CNS) (13)(14)(15). For example, in the CNS of the terrestrial slug *Limax*, electrophysiological studies have demonstrated the electrical synapses exist in the procerebral lobe (16) and in the identified neurons of the buccal ganglion (17). Recent studies have also performed the genomic and transcriptome analyses of various molluscan species (18)(19)(20)(21)(22). Whereas, there are limited studies on the molluscan innexin genes. In the pteropod mollusk *Clione limacina*, early studies identified two putative gap junction protein genes (23), and demonstrated that the electrical coupling was altered by the mRNA injection of one gap junction protein-coding gene (24). Recently, eight innexin gene transcripts were identified in the gastropod mollusk *Lymnaea stagnalis* (25). The sequence information of molluscan innexins is still limited and these identified innexin homologs have not been classified.

This study aimed to identify innexin gene transcripts, including circular RNAs (circRNAs), in the CNS of the terrestrial slug *Limax valentianus*. Additionally, comparative and phylogenetic analyses of the predicted amino acid sequences of molluscan species were performed to classify the innexin homologs.

## Methods

### Animals

All experiments were performed using the terrestrial slugs *L. valentianus* at 3–4 months post-hatching. The animals were maintained under laboratory conditions at 19 °C with a 12-h light/dark cycle. To anesthetize the animals before dissection, magnesium chloride solution was injected. The isolated CNS was frozen in liquid nitrogen for RNA extraction.

### Protein sequence analyses of predicted innexin homologs

Phylogenetic analysis was performed using the available molluscan innexin protein sequences in the Swissprot, Genbank, and Refseq databases with basic local alignment search tool (BLAST). Multiple sequence alignment of molluscan innexin homologs was performed using MUSCLE. The phylogenetic tree was constructed using the maximum likelihood method with MEGAX software (26).

The potential phosphorylation sites of protein kinase C (PKC), protein kinase A (PKA), protein kinase G (PKG), casein kinase 1 (CK1), casein kinase 2 (CK2), proto-oncogene tyrosine-protein kinase Src (SRC), P34cdc (cdc2), calmodulin-dependent protein kinase (CaMK), and mitogen-activated protein kinase (p38MAPK) were predicted using NetPhos 3.1 Server (27). Potential S-nitrosylation and N-glycosylation sites were predicted using GPS-SNO1.0 (28) and NetNGlyc 1.0 Server analysis (http://sno.biocuckoo.org/ and http://www.cbs.dtu.dk/services/NetNGlyc/), respectively. The consensus peptide sequence for N-glycosylation (Asn-X-Ser/Thr; X is not Pro) is conserved in both vertebrates and invertebrates(29)(30).

### Polymerase chain reaction (PCR)

The sequence-specific primers for *Limax* innexin mRNAs were designed based on the innexin homolog sequences in the *Limax* CNS identified through transcriptome shotgun assembly (TSA) (under published). For amplifying circRNA, divergent primers were designed based on the verified complementary DNA (cDNA) sequences. The primer sequences are listed in Supplemental Table 1. RNA was extracted from the dissected CNS of three animals using the Nucleospin RNA XS kit and treated with DNAse I (Macherey & Nagel, Düren, Germany). Total RNA (100 ng) was reverse-transcribed using M-MLV reverse transcriptase (Invitrogen, Carlsbad, CA) and random primers, following the manufacturer’s instructions. The cDNA was subjected to PCR using ExTaq DNA polymerase (Takara Co., Otsu, Japan) or PrimeSTAR GXL DNA polymerase kit (Takara Co.). The amplicons were subcloned into the TOPO™ vector (Invitrogen) and subjected to nucleotide sequencing analysis.

**Table 1.** Summary of the predicted modification sites in the innexin orthologs of the seven groups. The conserved modification sites between *Limax* innexin sequence and at least one other ortholog sequence are shown.

|  |  | PKC | PKA | CK1 | CK2 | SRC | cdc2 | N-glycosylation | S-nitrosylation |
| --- | --- | --- | --- | --- | --- | --- | --- | --- | --- |
| Group 1 | *Limax* Inx1 | S12, T170, S324, S338 | S131, S368 | S2 | S138, T374, S408 | - | - | - | - |
|  | *Limax* Inx2 | S12, T170, S338 | S131, S368 | S2, T135 | S138, T374, S408 | - | - | - | - |
| Group 2 | *Limax* Inx3 | T153, S337 | S363 | S369 | - | Y154 | - | N227 | - |
|  | *Limax* Inx4 | T155 | S365 | - | - | - | - | - | - |
| Group 3 | *Limax* Inx5 | S12, S145 | S187, T200, T387 | S145 | S18, S21 | - | - | - | - |
| Group 4 | *Limax* Inx6 | S13, T160, S178, T185, S341, S363, S388, S400 | S138, S144, S371, S392 | - | S381, S408 | Y151 | S135, S195, S324, S372 | - | C182 |
|  | *Limax* Inx7 | S7, T161, S172, T179, S185, S334, S357, S381, S383 | S128, S139, S364, S385 | - | S374, S401 | Y145 | S128, S318, S365 | - | C178 |
| Group 5 | *Limax* Inx8 | T34, S152, S182 | - | - | S359, S418 | - | - | - | C200, C212 |
|  | *Limax* Inx9x1 | T18, S133, S143 | - | - | S340, S369 | - | - | - | C181, C183 |
| Group 6 | *Limax* Inx10 | T173, T183, S355 | S200 | - | S152 | - | - | - | - |
| Group 7 | *Limax* Inx11x1 | T147, S160, T200, T326, S331, S334, S371 | S220 | S145, S180 | T147, T421 | - | - | N85, N102 | - |

## Results

### Identification of *Limax* innexin homologs

To identify the gap junction protein gene transcripts, a local BLASTX search was performed using *Aplysia* innexin homologs for TSA of *L. valentianus* CNS. The cDNA sequences from the TSA data were cloned and subjected to nucleotide sequencing. In this study, 16 *Limax* innexin homologs (*Limax* innexins 1–11, including spliced isoforms, and partial coding domain sequence (CDS) of *Limax* innexin 12: accession number LC595664–LC595679) were identified.

The deduced protein sequences were used for the prediction of transmembrane domain using OCTOPUS software (31). All obtained *Limax* innexin homologs exhibited typical characteristics of innexin proteins (four transmembrane domains that are connected by the first and second extracellular loops and one intracellular loop) (Fig. 1). Amino acid alignment analysis further revealed that the identified *Limax* innexins have a P-X-X-X-W motif around the second transmembrane domain (e.g. *Limax* innexin 1 PNIFW_120-124_; Fig. 1). The proline residue in this motif has been reported to function as a molecular hinge for voltage-dependent gating of connexin channels (32)(33). This motif is conserved in all the innexin/pannexin family proteins (34).

**Figure 1.**
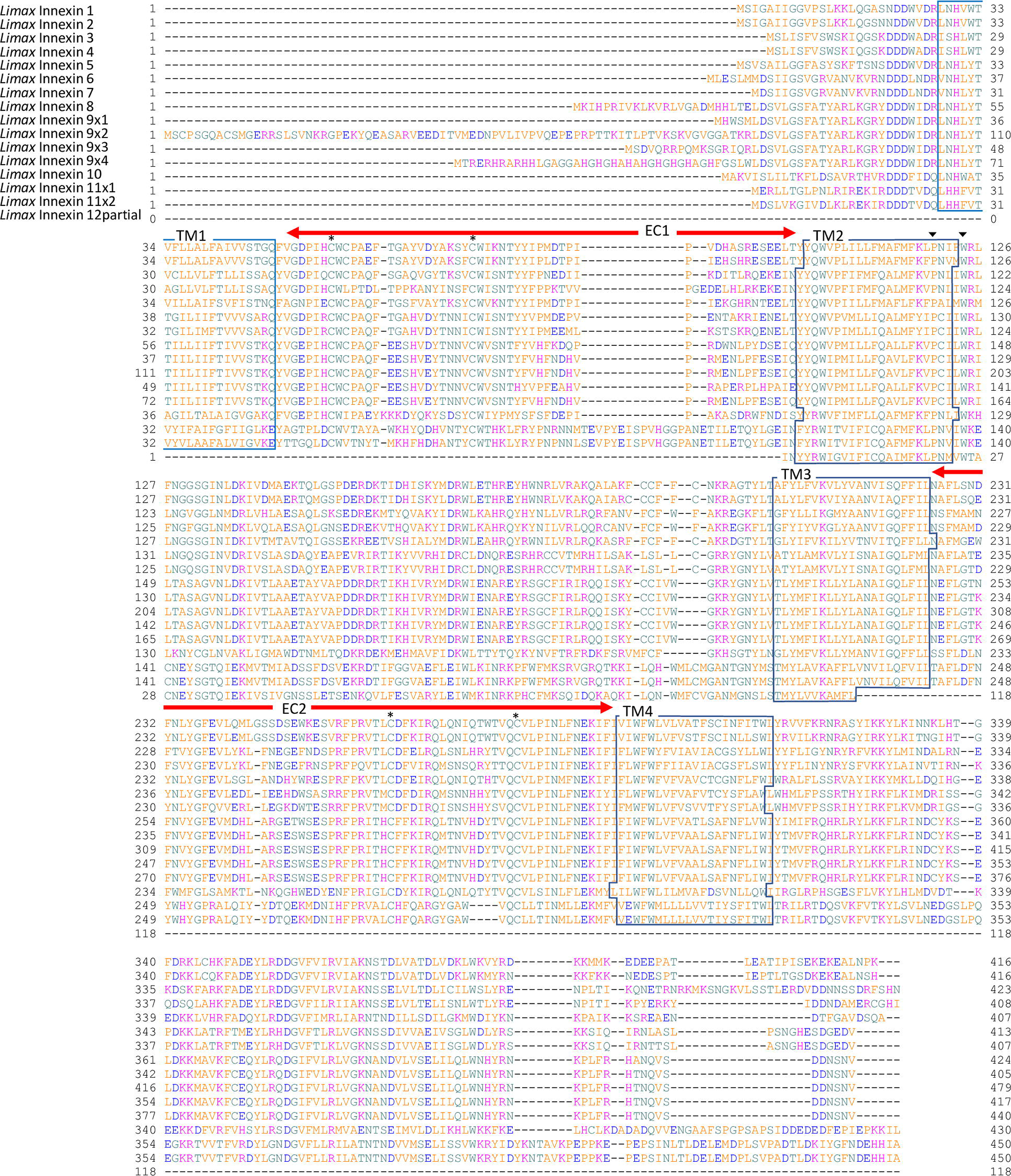
*Limax* innexin transcripts expressed in the central nervous system Alignment of deduced putative amino acid sequences of the *Limax* innexin homologs identified in the transcriptome of the central nervous system. The boxes show the four predicted transmembrane sites (TM1 to TM4), while the red arrows indicate two extracellular loops (EC1 and EC2). The asterisks indicate the extracellular cysteine residues in the extracellular loops, while the black arrow-heads indicate the P-X-X-X-W motif.

Another typical characteristic of innexin proteins is the conserved cysteine (Cys) residues in the extracellular loops (34). In invertebrate innexin, two pairs of Cys form essential intramolecular disulfide bonds between the first and second extracellular loops (35). The *Limax* innexin homologs have two Cys residues in the first and second extracellular loops (e.g. *Limax* innexin 1, Cys_56_ and Cys_74_ in the first extracellular loop; Cys_262_ and Cys_279_ in the second extracellular loop) (Fig. 1).

Interestingly, *Limax* innexins 1–9, except *Limax* innexin 4, have an additional Cys between the two Cys residues in the first extracellular loop (e.g. *Limax* innexin 1, Cys_58_) (Fig. 1). The additional Cys residue was also detected in the same region of other innexin/pannexin family protein sequences of some species listed in the public databases. In invertebrates, the additional Cys residue was detected in six of the 12 leech innexins (*Hirudo medicinalis* innexins 1, 3, 6, 9, 11, and 12). However, the additional Cys residue was not detected in the innexin sequences of insects and nematodes. In vertebrates, the additional Cys residue was detected in pannexin 3 of the clade *Cetartiodactyla* but not in pannexin 1 and 2, or connexin sequences (data not shown). These results indicate that the additional Cys residue is not a characteristic feature of molluscan innexins. However, the function of this additional Cys residue has not been reported in the innexin and pannexin proteins of any animal species.

The sequence alignment analysis also revealed differences in the identified *Limax* innexins (Fig. 1). The lengths of the extracellular loops were well conserved among *Limax* innexins 1–10 (the first extracellular loops, 53–55 amino acid residues; the second extracellular loop, 66–68 amino acid residues). However, only the length of the first extracellular loop in *Limax* innexin 11 was longer than that in other homologs (the first extracellular loop, 69 amino acids; the second extracellular loop, 63 amino acids). Furthermore, *Limax* innexins 1–10 had similar spacing between the two Cys residues in the extracellular loops. Meanwhile, the spacing between the two Cys residue in the second extracellular loop of *Limax* innexin 11 was short (*Limax* innexins 1–10, 16 amino acids; *Limax* innexins 11, 12 amino acids). Previous studies on vertebrate connexins have reported that the amino acid sequences between the Cys residues in the extracellular loop are critical for the formation of functional intercellular channels (9)(36)(37). Thus, we here suggest that *Limax* innexin isoforms can be broadly classified into two groups (*Limax* innexins 1–10 and *Limax* innexin 11). Interestingly, the present findings also appeared similar to the previous reports that vertebrate connexin and pannexin differ in the lengths of the extracellular loops (38)

### Molecular phylogenetic analysis of molluscan innexin homologs

Next, molecular phylogenetic analysis was performed to classify and examine the characteristics of identified *Limax* innexins. The innexin genes were reported to exhibit phylum-specific diversification in each phyla Arthropoda, Nematoda, and Mollusca (39). Hence, phylogenetic analysis was performed on the available molluscan innexin sequences in the Swissprot, Genbank, and Refseq protein databases. Additionally, the hypothetical protein sequences from the genomic data of *Elysia chlorotica* (BioProject PRJNA484060; sequence and assembled genome) and previously reported molluscan innexin sequences were used for the phylogenetic analysis (24)(25). In the pond snail *Lymnaea stagnalis*, in addition to the reported eight innexin transcripts (LstInx1 to LstInx8) (25), two homolog sequences were identified in another transcriptome data of the CNS (*Lymnaea* FX186689 and *Lymnaea* FX187001) (19). One variant of each gene was used if splicing variants were detected. The accession numbers for the same amino acid sequence are summarized in Figure 2 (e.g. *Crassostrea gigas* unc-9 isoformx1: EKC18227.1, XP011418374.1). The numbers of sequences in different animal species were as follows: *n*=20 for *Aplysia californica* (Gastropoda); *n*=10 for *Lymnaea stagnalis* (Gastropoda); *n*=2 for *Clione limacina* (Gastropoda); *n*=14 for *Biomphalaria glabrata* (Gastropoda); *n*=11 for *Pomacea canaliculata* (Gastropoda); *n*=12 for *Lottia gigantea* (Gastropoda); *n*=14 for *Elysia chlorotica* (Gastropoda); *n*=20 for *Mizuhopecten yessoensis* (Bivalvia); *n*=14 for *Crassostrea gigas* (Bivalvia); *n*=9 for *Octopus bimaculoides* (Cephalopoda); n=1 for *Sepioteuthis lessoniana* (Cephalopoda).

**Figure 2.**
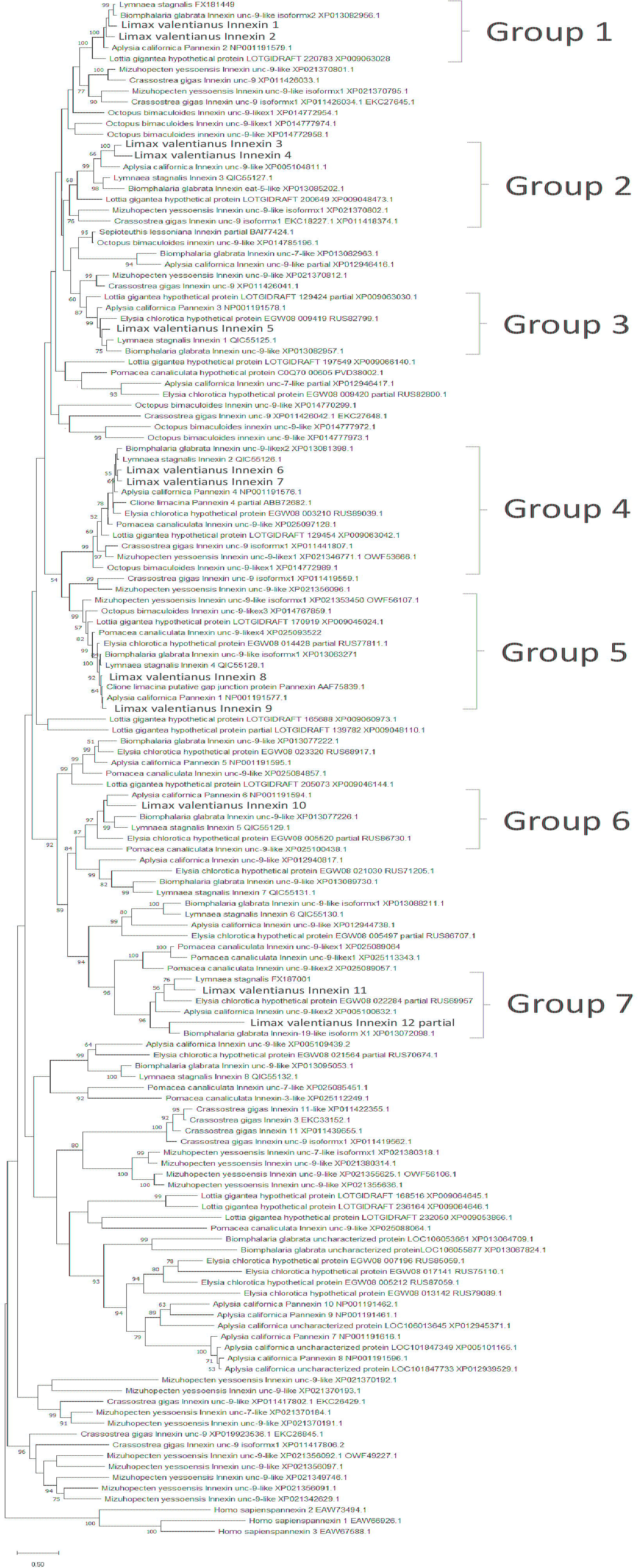
Phylogenetic tree of molluscan innexins Phylogenetic analyses of molluscan innexins were performed using the maximum likelihood method with *Homo sapiens* pannexins as the outgroup. The percentage of replicate trees in which the associated taxa clustered together in the bootstrap test (2000 replicates) is shown next to the branches (only values over 50 are displayed). Sequence accession numbers are indicated next to species names. The names of *Limax* innexin homologs are in bold font.

Molecular phylogenetic analysis revealed that the identified *Limax* innexin gene transcripts expressed in the CNS can be classified into seven ortholog groups (Fig. 2; Group 1, *Limax* innexins 1 and 2; Group 2, *Limax* innexins 3 and 4; Group 3, *Limax* innexin 5; Group 4, *Limax* innexins 6 and 7; Group 5, *Limax* innexins 8 and 9; Group 6, *Limax* innexin 10; Group 7, *Limax* innexins 11 and 12). The number of innexin homolog genes varied among species (e.g. *n*=20 for *Aplysia californica*; *n*=14 for *Elysia chlorotica*). Consistent with this finding, a previous study on vertebrate connexins demonstrated that the number of connexin genes in zebrafish (n=37) was approximately two times higher than that in humans (n=20) and mice (n=19) (40). The authors suggested that gene duplication events occur continuously in each phylum, which contribute to the species-specific regulation by gap junction-related proteins.

### Prediction of post-translational modification sites: N-glycosylation, S-nitrosylation, and phosphorylation

To classify the molluscan innexin homologs, potential protein modification sites in the deduced amino acid sequences of the seven groups were examined. The predicted modification sites in the innexin sequences of each group are shown in Figures 3–9. The conserved modification sites between orthologs are summarized in Figure 10.

**Figure 3.**
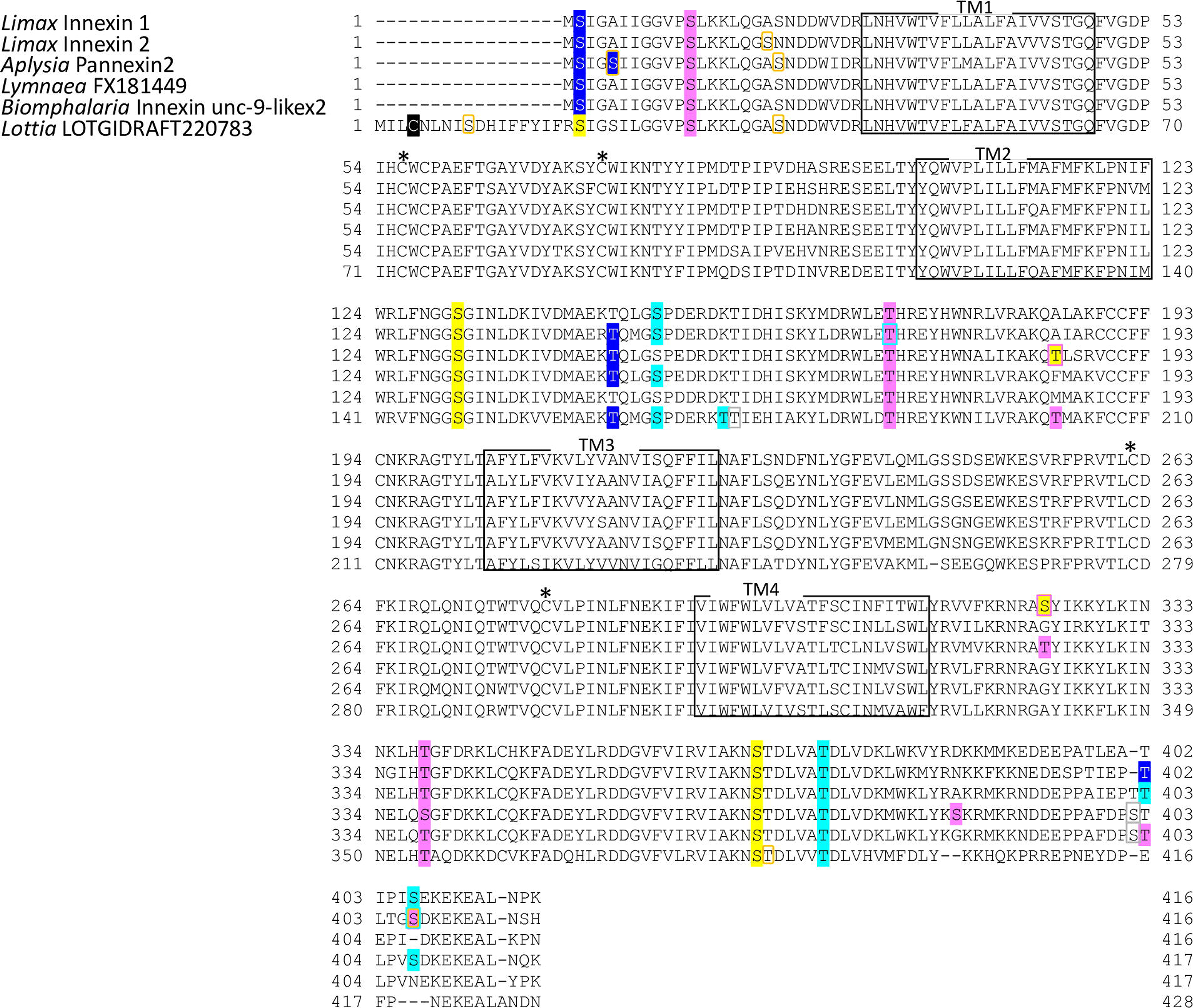
Protein sequence alignment of molluscan innexin orthologs in Group 1 Alignment of putative amino acid sequences of *Limax* innexin 1 and 2 with orthologous innexin sequences. The boxes show the four transmembrane sites, while the asterisks indicate the extracellular cysteine residues. The potential S-nitrosylation sites are shaded in black. The phosphorylation sites are indicated with shading or box in magenta (protein kinase C), yellow (protein kinase A), blue (casein kinase 1), cyan (casein kinase 2), orange (P34cdc), and gray (protein kinase G).

**Figure 4.**
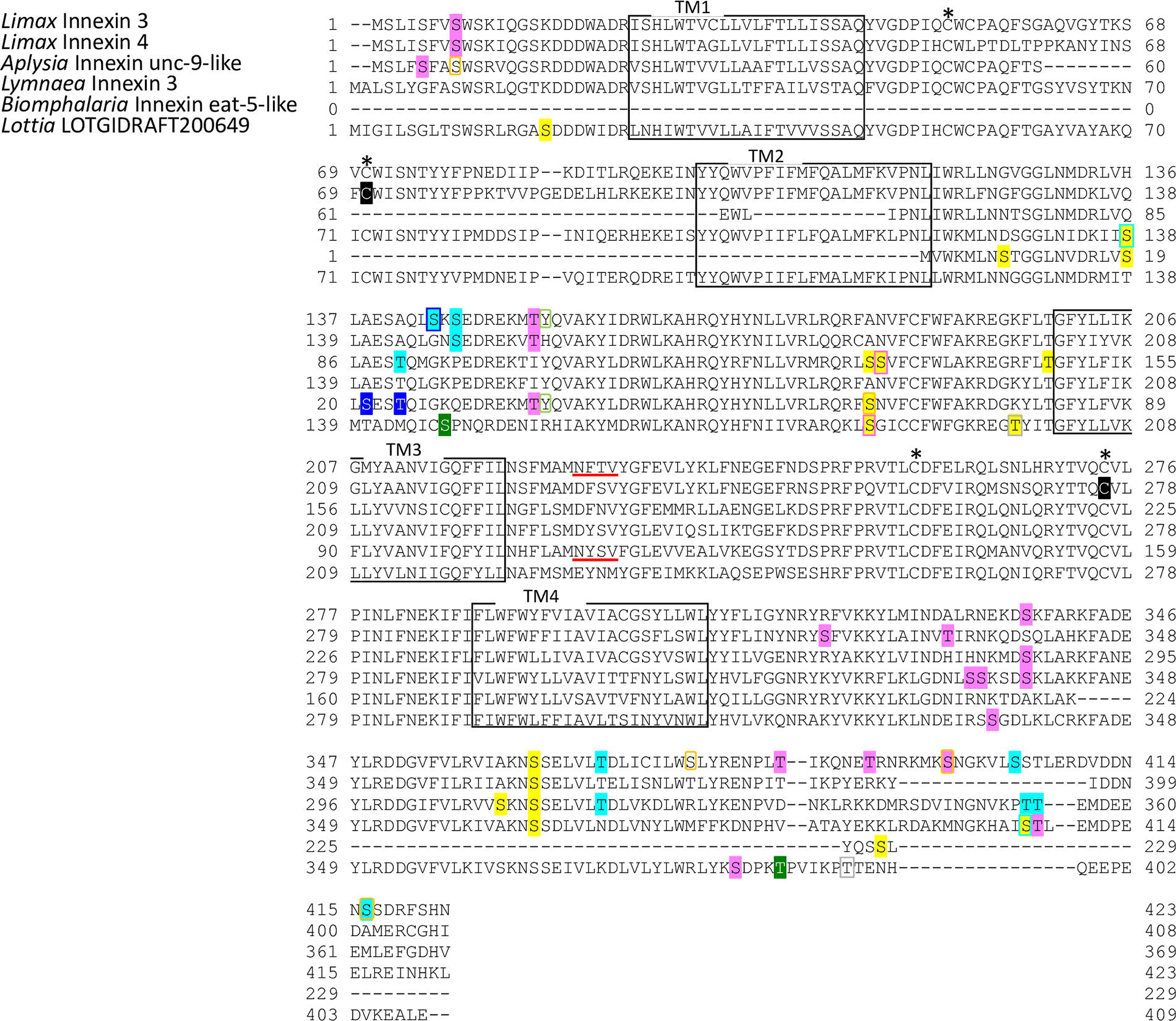
Protein sequence alignment of molluscan innexin orthologs in Group 2 Alignment of putative amino acid sequences of *Limax* innexin 3, 4 with orthologous innexin sequences. The boxes show the four transmembrane sites, while the asterisks indicate the extracellular cysteine residues. The potential S-nitrosylation sites are shaded in black. The phosphorylation sites are indicated with shading or box in magenta (protein kinase C), yellow (protein kinase A), blue (casein kinase 1), cyan (casein kinase 2), orange (P34cdc2), green (p38-mitogen-activated protein kinase), light green (proto-oncogene tyrosine-protein kinase Src), and gray (protein kinase G). Potential N-glycosylation sites are underlined with bold red lines.

**Figure 5.**
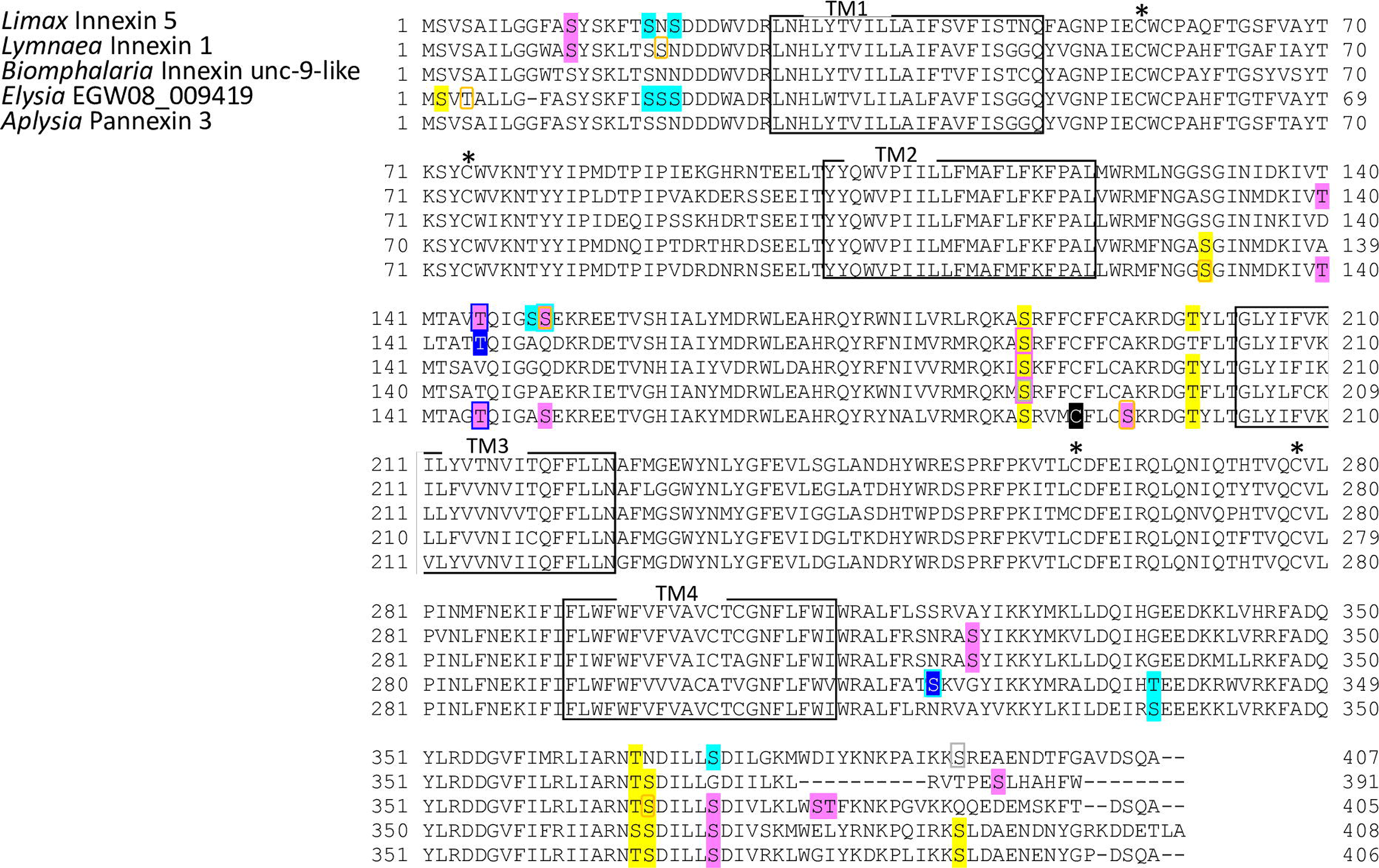
Protein sequence alignment of molluscan innexin orthologs in Group 3 Alignment of putative amino acid sequences of *Limax* innexin 5 with orthologous innexin sequences. The boxes show the four transmembrane sites, while the asterisks indicate the extracellular cysteine residues. The potential S-nitrosylation sites are shaded in black. The phosphorylation sites are indicated with shading or box in magenta (protein kinase C), yellow (protein kinase A), blue (casein kinase 1), cyan (casein kinase 2), orange (P34cdc2), and gray (protein kinase G).

Potential N-glycosylation sites were detected only in the innexin homologs of Groups 2 and 7. In Group 2, one potential N-glycosylation site was identified at the first extracellular loop in two of the six orthologs (*Limax* innexin 3, N_227_FT; *Biomphalaria* XP013085202.1 partial CDS sequence, N_110_YS). In Group 7, two N-glycosylation sites were identified in the first extracellular loop in all orthologs of Group 7 (*Limax* innexin 11 splice variants, *Limax* innexin 11×1, N_85_MT and N_102_ET; *Limax* innexin 11×2, N_85_LS and N_102_ET; *Lymnaea* FX187001, N_85_LS and N_102_ET; *Elysia* EGW08_022284 partial sequence, N_102_AT; *Biomphalaria* innexin-19-likex1 N-terminal partial sequence, N_118_LS and N_134_TT; *Aplysia* innexin unc-9-likex2, N_85_LS and N_101_LT), except *Limax* innexin 12 for which the sequence information of the whole coding region was not available.

This indicated that N-glycosylation modification occurs in the molluscan innexins of certain ortholog groups. The N-glycosylation sites of molluscan innexins are conserved in the first extracellular loop of all orthologs in Group 7. Consistent with this finding, vertebrate pannexin proteins have N-glycosylation sites at the extracellular loops, whereas almost all connexins do not have the N-glycosylation sites.

N-glycosylation of pannexins regulates subcellular localization and intermixing of different pannexins for channel formation (41)(42). Additionally, N-glycosylation of pannexin inhibits the intercellular docking of neighboring pannexin channels and the formation of gap junctions (43)(42). Interestingly, molluscan innexin sequences of Group 7 exhibited the following two pannexin-like characteristics: N-glycosylation at the extracellular loops and longer extracellular loops than the innexin homologs in other groups.

Next, the S-nitrosylation sites were examined in the molluscan innexin homologs of the seven groups. The S-nitrosylation sites were identified in some homologs. The conserved S-nitrosylation sites were identified only in the intracellular loops of the orthologs of Groups 4 and 5 (Figs. 6, 7 and 10). S-nitrosylation, which is the covalent addition of nitric oxide (NO) to the Cys residue, is reported to be involved in the regulation of channel properties, such as channel permeability and voltage-sensitive gating of vertebrate connexins and pannexins (44)(45)(46). Various studies have examined the role of NO in the CNS of molluscan species (47)(48)(49)(50)(17). The findings of this study indicate that NO may regulate some molluscan innexins that belong to phylogenetically related groups.

**Figure 6.**
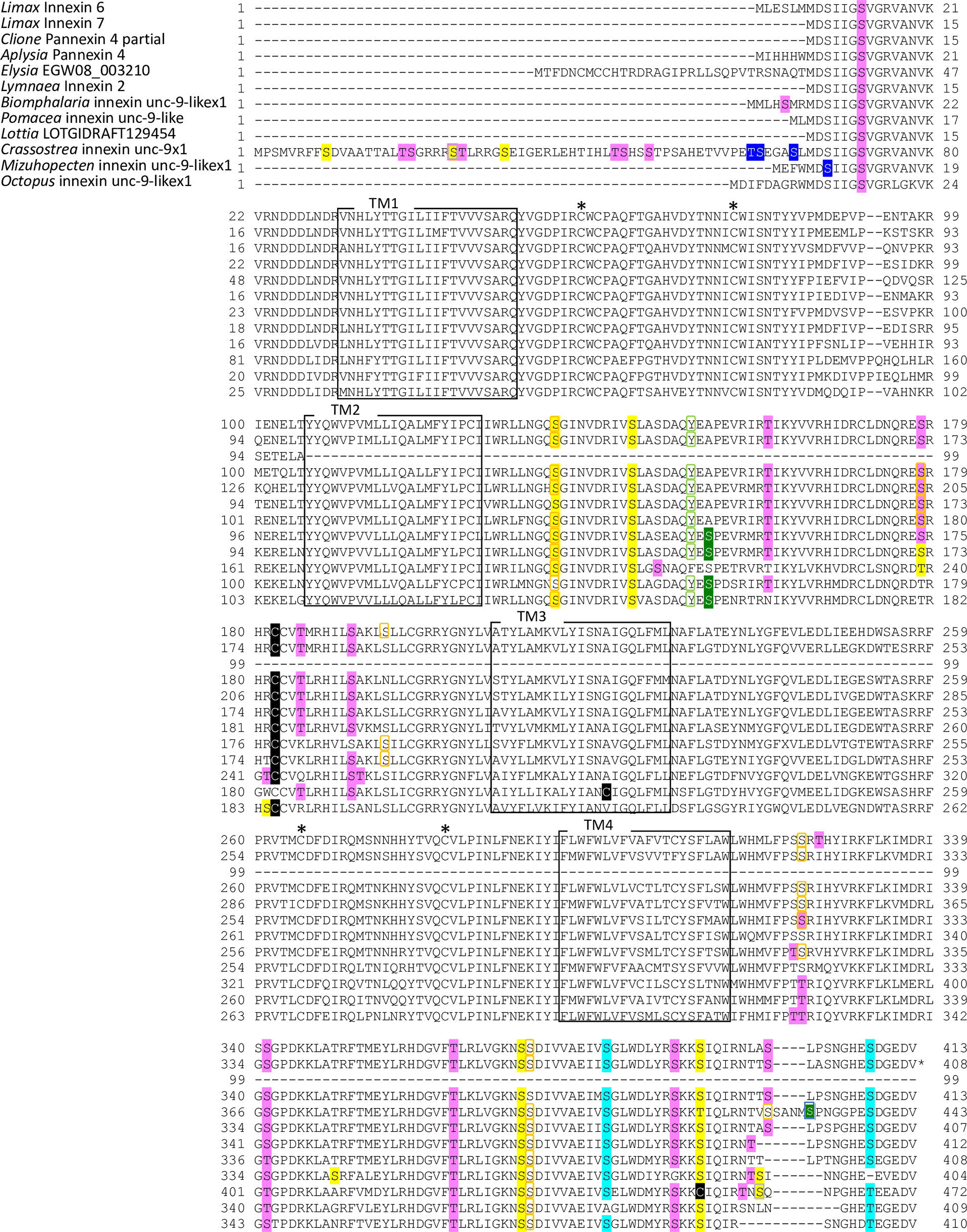
Protein sequence alignment of molluscan innexin orthologs in Group 4 Alignment of putative amino acid sequences of *Limax* innexins 6 and 7 with orthologous innexin sequences. The boxes show the four transmembrane sites, while the asterisks indicate the extracellular cysteine residues. The phosphorylation sites are indicated with shading or box in magenta (protein kinase C), yellow (protein kinase A), blue (casein kinase 1), cyan (casein kinase 2), orange (P34cdc2), green (p38-mitogen-activated protein kinase), light green (proto-oncogene tyrosine-protein kinase Src), and gray (protein kinase G).

**Figure 7.**
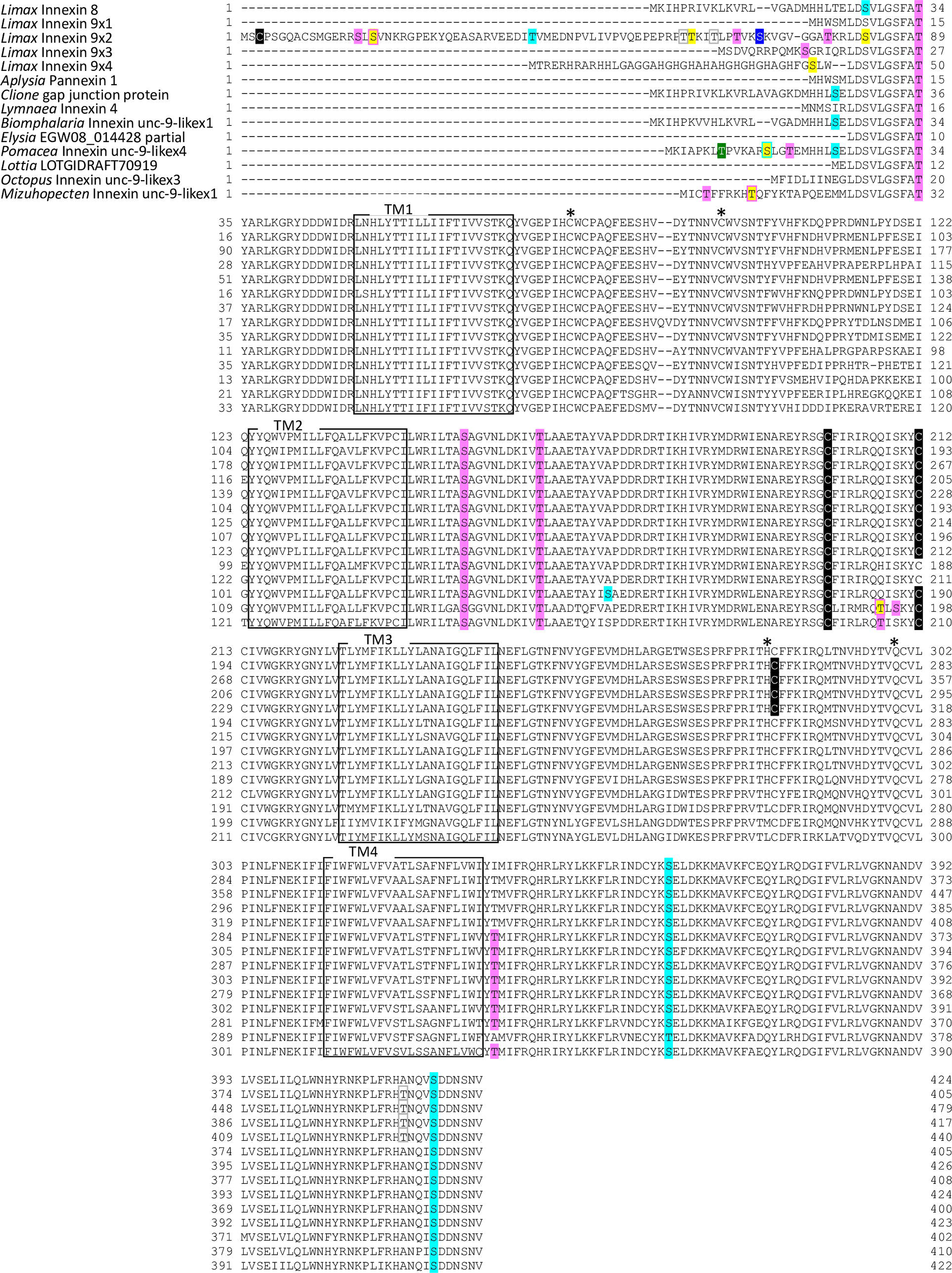
Protein sequence alignment of molluscan innexin orthologs in Group 5 Alignment of putative amino acid sequences of *Limax* innexins 8 and 9 with orthologous innexin sequences. The boxes show the four transmembrane sites, while the asterisks indicate the extracellular cysteine residues. The potential S-nitrosylation sites are shaded in black. The phosphorylation sites are indicated with shading or box in magenta (protein kinase C), yellow (protein kinase A), blue (casein kinase 1), cyan (casein kinase 2), green (p38-mitogen-activated protein kinase), and gray (protein kinase G). Potential N-glycosylation sites are underlined with bold red lines.

**Figure 8.**
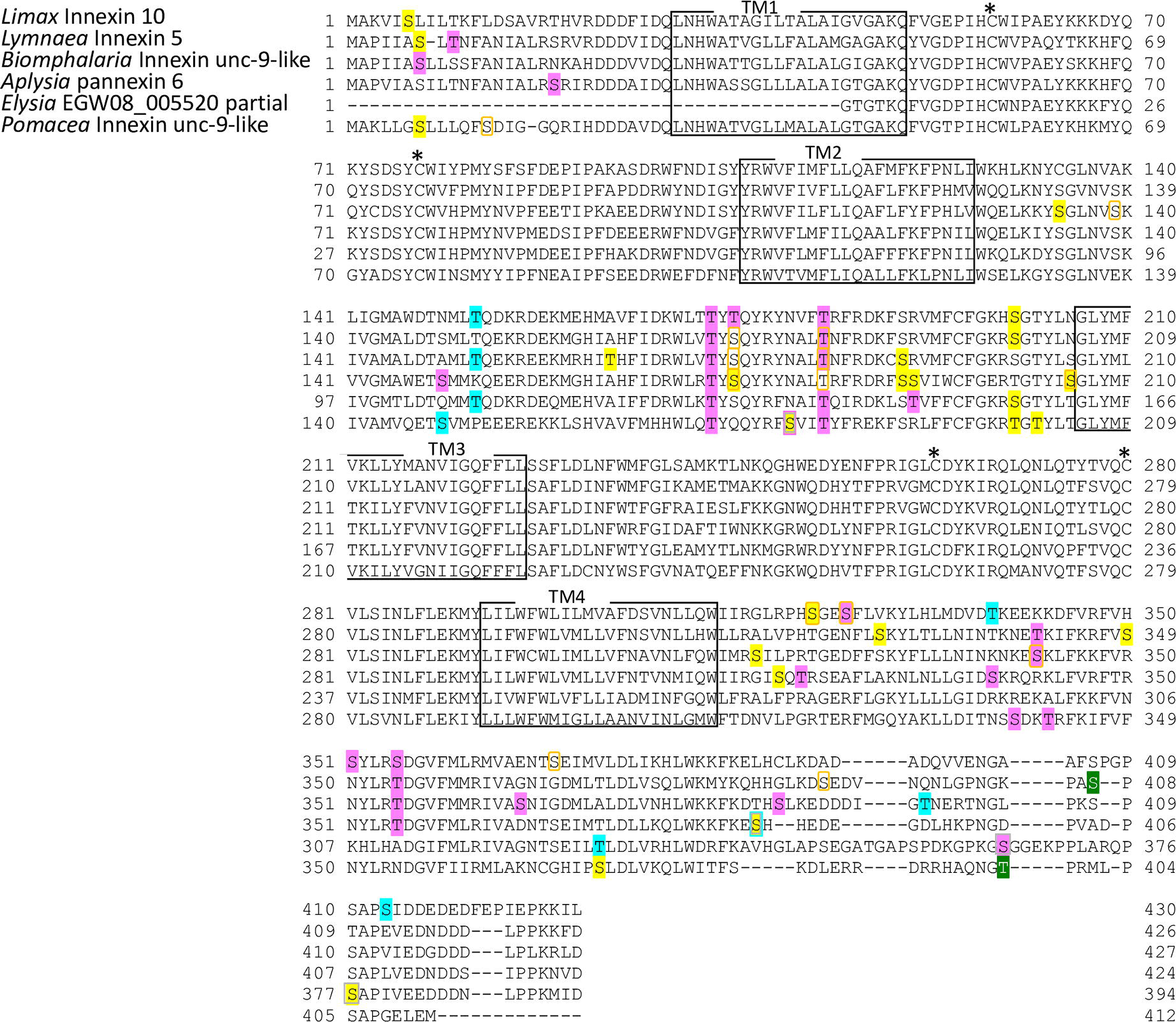
Protein sequence alignment of molluscan innexin orthologs in Group 6 Alignment of putative amino acid sequences of *Limax* innexin 10 with orthologous innexin sequences. The boxes show the four transmembrane sites, while the asterisks indicate the extracellular cysteine residues. The phosphorylation sites are indicated with shading or box in magenta (protein kinase C), yellow (protein kinase A), blue (casein kinase 1), cyan (casein kinase 2), orange (P34cdc2), green (p38-mitogen-activated protein kinase), and gray (protein kinase G).

**Figure 9.**
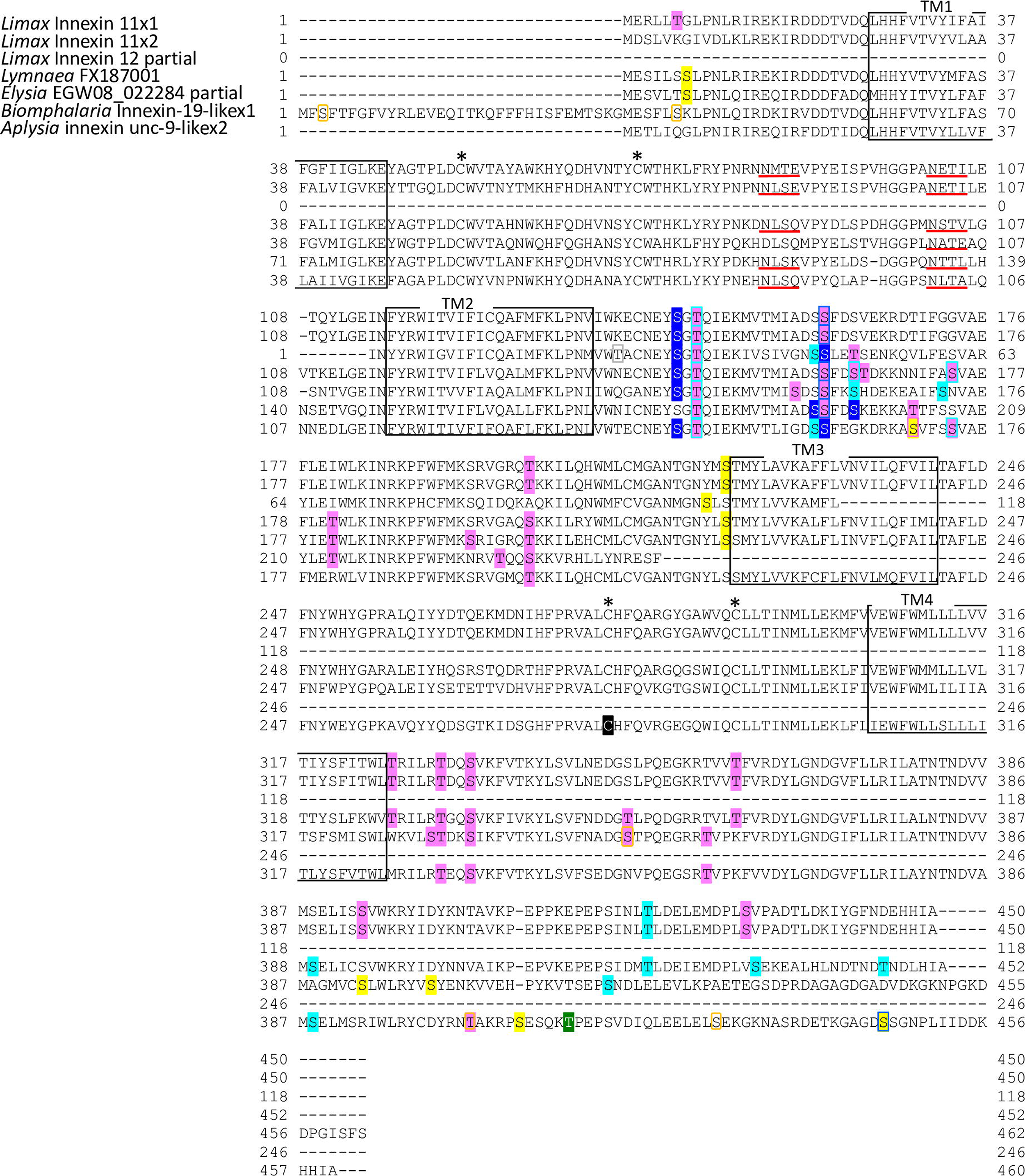
Protein sequence alignment of molluscan innexin orthologs in Group 7 Alignment of putative amino acid sequences of *Limax* innexins 11 and 12 with orthologous innexin sequences. The boxes show the four transmembrane sites, while the asterisks indicate the extracellular cysteine residues. The potential S-nitrosylation sites are shaded in black. The phosphorylation sites are indicated with shading or box in magenta (protein kinase C), yellow (protein kinase A), blue (casein kinase 1), cyan (casein kinase 2), orange (P34cdc2), green (p38-mitogen-activated protein kinase), and gray (protein kinase G). Potential N-glycosylation sites are underlined with bold red lines.

Further, the phosphorylation sites of several kinases, including PKC, PKA, PKG, CK1, CK2, SRC, cdc2, CaMK, and p38MAPK, were examined (Figs. 3–9). These kinases phosphorylate vertebrate connexin/pannexin and regulate protein functions, such as oligomerization, channel properties, intracellular trafficking, gap junction assembly, and stability (5)(51)(6)(52). The characteristics of the sequences in each ortholog group are shown in Figures 3–9 and 10. The phosphorylation sites in PKG, CaMK, and p38MAPK were not conserved in the ortholog groups. Among the seven ortholog groups, the number of phosphorylation sites was high in Group 4. The conserved phosphorylation sites of cdc2 were detected only in this group (Figs. 6 and 10). Although Groups 4 and 5 were phylogenetically related (Fig. 2), the innexin homologs of Group 5 exhibited a decreased number of phosphorylation sites (Figs. 7 and 10). This suggested that the predicted phosphorylation sites are conserved among the orthologs in each group and that each group exhibits a distinctive phosphorylation pattern.

Among the seven groups, the orthologs of Group 2 exhibited low sequence homology. Additionally, the protein modification sites in the orthologs of Group 2 were more variable than those in the orthologs of other groups (Figs. 4 and 10). Interestingly, *Limax* innexins 3 and 4 exhibited the highest degree of sequence homology. However, the protein modification sites were different between *Limax* innexins 3 and 4 (Figs. 2 and 10). At the extracellular loop, *Limax* innexin 3 had an N-glycosylation site, while *Limax* innexin 4 had two S-nitrosylation sites. The number of phosphorylation sites in the intracellular carboxyl-terminal domain of *Limax* innexin 3 was higher than that in the intracellular carboxyl-terminal domain of *Limax* innexin 4 (T_385_, T_391_, and S_398_ for PKC; T_369_, S_405_, and S_416_ for CK2; and S_377_ and S_398_ for cdc2). The difference between molluscan innexin genes in Group 2 could arise from continuous duplication events, which may result in new functions for the innexin proteins in Group 2.

### CircRNA identification of *Limax* innexin genes

Next, RT-PCR experiments were performed to examine the expression of *Limax* innexin transcripts in the CNS. Interestingly, multiple amplicons were obtained for some innexin homologs. Sequencing analysis revealed that these amplicons were circRNAs. CircRNA is a class of non-coding RNAs that are generated through alternative back-splicing, which connects the terminal 5’ and 3’ ends of linear pre-mRNA (53). Previous studies have demonstrated the presence of circRNAs in different animal species, such as mammals, insects, and nematodes (54)(55).

Interestingly, circRNA is abundant in the brain tissue and the circRNA-producing genes encode proteins related to synapse-related functions (54)(56). However, the function of circRNA has not been fully elucidated.

Previous studies indicate that circRNAs mainly comprise coding exons (55). To identify the circRNAs of the innexin genes, divergent primers were designed for the coding region of *Limax* innexin sequences. This study identified 16 circRNA sequences for nine *Limax* innexin genes (Table 2). CircRNA formation is observed among *Limax* innexins in the CNS. Some circRNAs encode the whole coding region (circINX2a, circINX3, and circINX7b). The result demonstrated that circRNAs are constitutively formed from the innexin genes in the CNS of *Limax*.

**Table 2.**
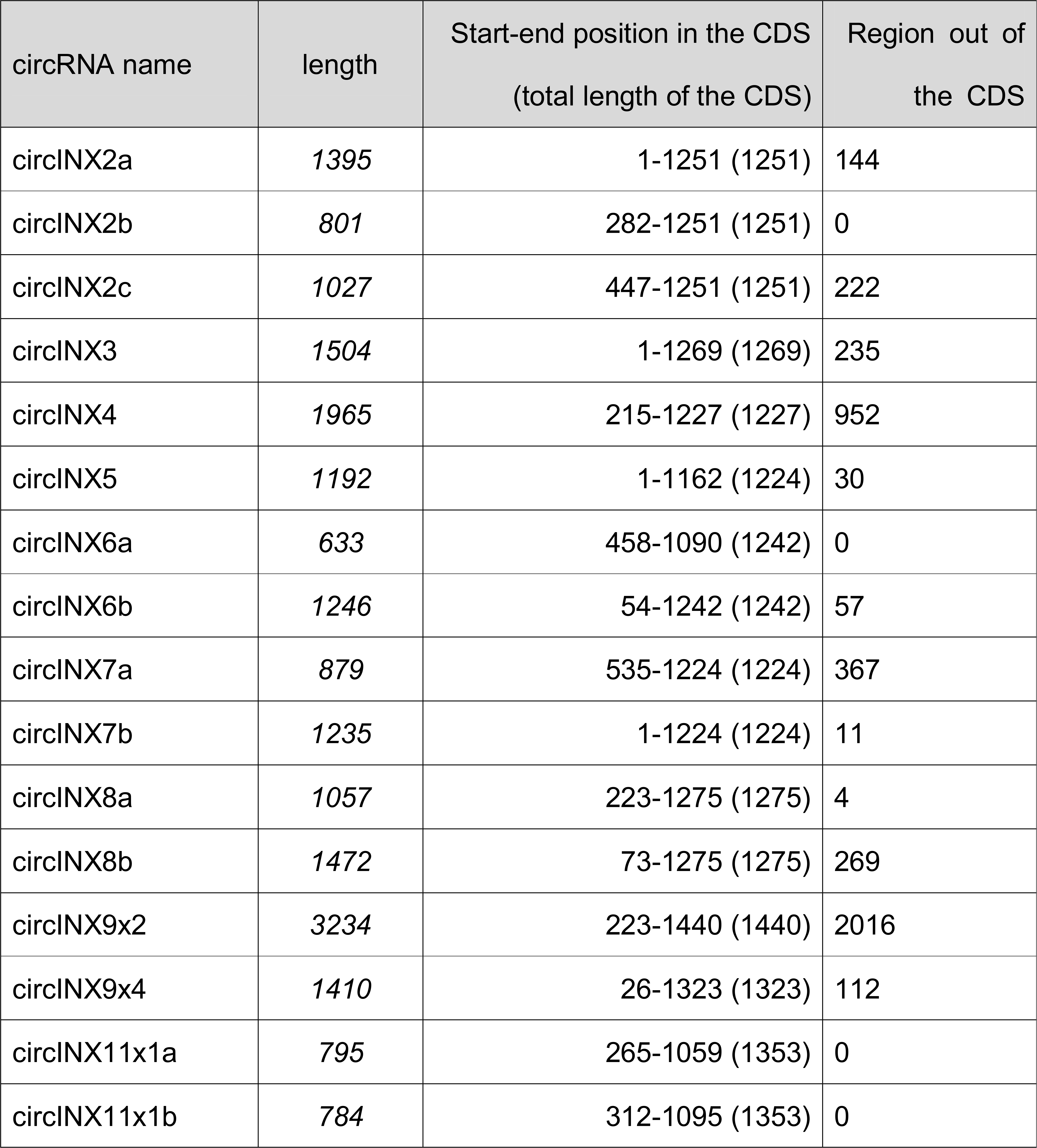
Summary of identified *Limax* innexin circular RNAs (circRNAs) in the central nervous system. CDS, coding domain sequence.

| circRNA name | length | Start-end position in the CDS<br>(total length of the CDS) | Region out of<br>the CDS |
| --- | --- | --- | --- |
| circINX2a | 1395 | 1-1251 (1251) | 144 |
| circINX2b | 801 | 282-1251 (1251) | 0 |
| circINX2c | 1027 | 447-1251 (1251) | 222 |
| circINX3 | 1504 | 1-1269 (1269) | 235 |
| circINX4 | 1965 | 215-1227 (1227) | 952 |
| circINX5 | 1192 | 1-1162 (1224) | 30 |
| circINX6a | 633 | 458-1090 (1242) | 0 |
| circINX6b | 1246 | 54-1242 (1242) | 57 |
| circINX7a | 879 | 535-1224 (1224) | 367 |
| circINX7b | 1235 | 1-1224 (1224) | 11 |
| circINX8a | 1057 | 223-1275 (1275) | 4 |
| circINX8b | 1472 | 73-1275 (1275) | 269 |
| circINX9x2 | 3234 | 223-1440 (1440) | 2016 |
| circINX9x4 | 1410 | 26-1323 (1323) | 112 |
| circINX11x1a | 795 | 265-1059 (1353) | 0 |
| circINX11x1b | 784 | 312-1095 (1353) | 0 |

The functions of circRNAs generated from the connexin, pannexin, and innexin genes have not been elucidated. Thus, the circRNAs of vertebrate connexin and pannexin genes were examined using a database that contains >32,000 human circRNAs (57). CircRNA data were retrieved for four connexin genes (GJA1, connexin 43; GJB3, connexin 31; GJB5, connexin 31.1; GJC1, connexin 45) and one pannexin gene (PNX1, pannexin 1). Of these, circRNAs of connexin 45 (hsa_circ_0106948, hsa_circ_0106949, hsa_circ_0106950, hsa_circ_0106951, hsa_circ_0106952) and pannexin 1 (hsa_circ_0096811, hsa_circ_0096812, hsa_circ_0096813, and hsa_circ_0096814) were listed as the transcripts in the brain tissue. These data suggest that circRNA formation from innexin genes is not a characteristic event in mollusks or invertebrates and that circRNAs of some homolog genes (not all genes) are involved in the regulatory mechanism of a single-membrane channel and gap junction.

## Discussion

In this study, the innexin homologs expressed in the CNS of *L. valentianus* were identified and characterized. Phylogenetic analysis revealed that innexin genes in molluscan species and the ortholog groups exhibited diversity. The prediction of post-translational modifications, including phosphorylation, S-nitrosylation, and N-glycosylation, revealed the characteristics of each group. Additionally, this study demonstrated the abundant expression of circRNA of innexin homologs in the CNS.

In invertebrate species, innexin is the only protein family and the multiple isoforms have not been classified. Based on the phylogenetic analysis, this study classified *Limax* innexin homologs in the CNS into seven groups of molluscan innexin orthologs. Interestingly, one ortholog group exhibited vertebrate pannexin-like characteristics with different lengths of the extracellular loops and glycosylation sites. Previous studies have demonstrated that the intercellular docking of two single-cell membrane channels is regulated by the interaction between the extracellular loops of gap junction-related proteins. Pannexin proteins mainly form single-cell membrane channels *in vivo*, and glycosylation of pannexin at the extracellular loop may inhibit the formation of intercellular channels (43) (12). So far, only one study has examined the glycosylation of innexin homologs in the yellow fever mosquito *Aedes aegypti* (58). The study reported that two of the six *Aedes* innexins (AeInx3 and AeInx7) have putative glycosylation sites and that AeInx3 exhibited multiple bands in the western blot. The authors suggested that AeInx3 may be an important component of gap junctions. However, the study did not discuss the effect of glycosylation on intercellular channel formation. Regard to the previous studies of pannexin, the glycosylation status of AeInx3 protein could regulate the formation of single-membrane channels or intercellular gap junction channels. In this study, in addition to innexin orthologs in Group 7, *Limax* innexin 3 in Group 2 had an N-glycosylation site, which is not conserved among the orthologs, at the extracellular loop. Thus, *Limax* innexin 3 could form both gap junction channels and single-cell membrane channels, and the N-glycosylation status of this protein could regulate the intercellular docking of the channels.

In addition to the N-glycosylation sites, this study predicted the phosphorylation and S-nitrosylation sites and characterized molluscan innexins of the seven groups. Each group exhibited a distinctive protein modification pattern. Additionally, the protein modification sites were well conserved among the homologs of the same group (Table 1). This indicated that the regulatory mechanism of the channels is well conserved among orthologs. These results also indicate that similar to that in vertebrate connexins (40), the duplication divergence of innexin genes could generate multiple innexin isoforms in molluscan species.

Further, this study examined the formation of circRNAs from innexin genes. This study also detected connexin and pannexin circRNAs in the human circRNA database (59). However, there is no study that experimentally examined the existence of circRNA for gap junction-related proteins. So far, only one review has proposed that mammalian connexin 43-derived circRNA functions as a miRNA sponge during breast cancer initiation stages (60). However, this has not been experimentally demonstrated. And the function of circRNAs is currently under investigation. Some studies have reported that circRNAs function as miRNA sponges (55)(61). Recent studies have reported that several circRNAs are associated with polysomes and that circRNAs are translated (62)(63). In this study, 12 of the 16 *Limax* innexin circRNAs encoded an open reading frame. These circRNAs can act as a template for N-terminal truncated innexin proteins. Interestingly, several studies have reported the unexpected functions of N-terminal truncated connexin that are generated from internal translation initiation (64)(65)(66). The short connexin 43 isoform is involved in trafficking the full-length connexin 43 to the gap junction site. Additionally, the short connexin 43 isoform directly regulates N-cadherin transcription, which results in neural crest cell migration at an early developmental stage in the amphibian and mammalian cells. The translation of innexin circRNA may result in the generation of truncated innexin proteins, which may have uncharacterized roles, such as protein trafficking and regulation of gene expression. To examine this hypothesis, future studies must examine the expression and function of innexin proteins.

## Conclusion

In this study, multiple innexin transcripts were identified in *L. valentianus* and the molluscan innexin sequences were phylogenetically classified. Additionally, circRNA formation was demonstrated in multiple innexin genes in the CNS, which provided novel insights for understanding the regulatory mechanism of gap-junction-related proteins. The findings of this study will facilitate further studies on the function of molluscan innexin proteins and contribute to future studies on the regulatory mechanisms of gap junction-related proteins.

## Acknowledgments

The authors would like to thank Tokushima Bunri University for financial support. The authors also thank Miyu Nakagawa, Satsuki Maejima, and Ko Sudo for helping with the molecular biology experiments.

## Notes

### Competing Interest Statement

The authors have declared no competing interest.

