## Supplemental Table 1 for "Identification and classification of innexin gene transcripts in the central nervous system of the terrestrial slug *Limax valentianus*"

PCR primers used in this study

| **Targeted mRNA or circRNA** | **Forward primer** | **Reverse primer** |
| --- | --- | --- |
| INX1 | CAACGAACCAAGGGATTTAAACTAGCGATT | CAAAGGTTAAAACACCTCCCATTTGCTC |
| INX2 | CGCTGTGAGACTCTCGAATGTCTATAGGG | CCTGAAAGTTGTGGAAGCGAAATGATTTTCTGTC |
| INX3 | AAAGAAATCCGGTGCGGTCTCA | TTTGGTTCACTTCATGCCCCAAA |
| INX4 | GCTATTTCGACGACTCGCCAGCTAAC | TCCAGGCCTGCCTACAACTCTACTC |
| INX5 | CTGTCCTCGTGCCGTTATTGAGACTTG | CCATAAAGACATGCCATTCAAGCTTGAGAG |
| INX6 | GAACCATGAGATGGACAGTATCATTGGGT | CTATACATCTTCGCCATCACTCTCATGGC |
| INX7 | AGTGAGACTCCAAATACAGCCCGAAGA | ATTCGATGAGGCTGTTCTAATGTCAAGGT |
| INX8 | GCCAATCAAAGGCTGAGATTTTCCGT | GGCTTGAAGACTTTATACCAAGCCAGTT |
| INX9x1 | GATTTGAACCTCTACAATCGGAGAGGAGT | ATGGGTCTCTTCCGGGGTAGGT |
| INX9x2 | AGCCTTAAGATACCGCGACCAGTT | ATGGGTCTCTTCCGGGGTAGGT |
| INX9x3 | CCCTGTCGTTGTGATAAATACTACAACTTCCTG | ATGGGTCTCTTCCGGGGTAGGT |
| INX9x4 | ACCCTGACAGGAGGTCTACTATCGTG | ATGGGTCTCTTCCGGGGTAGGT |
| INX10 | GCGCACAGAGATTTGTTATTTTGAAGAAGCGT | ACAGTGAACAATCACTGATTTGCCCAGTT |
| INX11x1 | GAAGTTTATTAGACAAATATGGAGCGGTTGCTGAC | GTTCTATCAGCTGCTAACCTTTTCTTAGTTGACCT |
| INX11x2 | AATAACTAGGTTGTCATGGATTCATTGGTTAAGG | CAATTCAATGTCCTGTTCTATCAGCTGCTAAC |
| circINX01 | TTGTACCGTGTTGTCTTCAAGAGGAACA | AGCCTCCGTTGAAGAGTCTCCAGAA |
| circINX02 | ACCTCAAGATCACCAACGGGATCCA | TGCATAGATCACCTTGACGAACAGGTAGAG |
| circINX03 | TCTCACAGATCTGATTTGCATTCTCTGGAGT | AACGTTTGCAAATCTCTGTCGGAGTCTG |
| circINX04 | AATTCCTCAGAATTAGTTCTCACGGAGCTCATC | GAAATTTACCCTCACGTTTGGCAAACCAG |
| circINX05 | GGGGAGTGGTACAATTTGTATGGGTTTGAAGTG | GAACTTGGAGTAGGATGCAAAGCCTCCTAGA |
| circINX06 | CGTTGCATTTGTAACCTGCTACAGTTTCC | ATAAACAACTGACCAATGGCGTTTGAGATG |
| circINX07 | CATGTGGTTCTGGCTGGTGTTCGTGT | GTAAGTGGCAACAAGATAGTTCCCATAGCGT |
| circINX08 | TGGAACCATTATAGAAACAAGCCCCTGTTC | TCGATCCATGTATCGGACAATGTGCT |
| circINX09 | TTAGACACACCAATCAAGTGAGCGATGACAAC | GATCCCGATCGTCTGGGGCTACATAGG |
| circINX10_1 | AAATTAGTGCGAGTGAAATCTGTGGAGTAG | GACCTTGCTTGTTCAAGGTCTTCATGG |
| circINX10_2 | AGATTGAGCCGCGGAGTTAGAGTG | GACCTTGCTTGTTCAAGGTCTTCATGG |
| circINX11x1_1 | CCAGTCCGTCAAGTTTGTTACGAAGTATCTCTC | TCACATTCGGCAGCTTGAACATGAAGG |
| circINX11x1_2 | ATGTCCACCATGTACCTTGCAGTCAAGGCT | GCCTGGCATATAAATATCACGGTGATCCA |
